# Measuring epistasis in fitness landscapes: the correlation of fitness effects of mutations

**DOI:** 10.1101/042010

**Authors:** Luca Ferretti, Benjamin Schmiegelt, Daniel Weinreich, Atsushi Yamauchi, Yutaka Kobayashi, Fumio Tajima, Guillaume Achaz

**Affiliations:** Evolution Paris-Seine (UMR 7138) and Atelier de Bio-Informatique, UPMC, Paris; SMILE, CIRB (UMR 7241); Collège de France, Paris, France.; The Pirbright Institute, Woking, United Kingdom.; Institute for Theoretical Physics, University of Cologne, Germany.; Ecology and Evolutionary Biology, Brown University, Providence, USA.; Center for Ecological Research, Kyoto University, Japan.; Kochi University of Technology, Japan.; Department of Biological Sciences, University of Tokyo, Japan.

**Keywords:** epistasis, fitness, mutations, ruggedness

## Abstract

Genotypic fitness landscapes are constructed by assessing the fitness of all possible combinations of a given number of mutations. In the last years, several experimental fitness landscapes have been completely resolved. As fitness landscapes are high-dimensional, simple measures of their structure are used as statistics in empirical applications. Epistasis is one of the most relevant features of fitness landscapes. Here we propose a new natural measure of the amount of epistasis based on the correlation of fitness effects of mutations. This measure has a natural interpretation, captures well the interaction between mutations and can be obtained analytically for most landscape models. We discuss how this measure is related to previous measures of epistasis (number of peaks, roughness/slope, fraction of sign epistasis, Fourier-Walsh spectrum) and how it can be easily extended to landscapes with missing data or with fitness ranks only. Furthermore, the dependence of the correlation of fitness effects on mutational distance contains interesting information about the patterns of epistasis. This dependence can be used to uncover the amount and nature of epistatic interactions in a landscape or to discriminate between different landscape models.

## 1. Introduction

Fitness landscapes represent a successful metaphor to understand evolution as a hill-climbing process. The seminal work of Sewall Wright [1] inspired a vast amount of theoretical work in phenotypic and molecular evolution [2, 3]. Furthermore, this metaphor contributed to the scientific exchange with other fields, especially with computer science [4] and physics [5]. Recently, Wright’s idea of genotype-fitness landscapes moved from a metaphor to an object of experimental studies, as several fitness landscapes were experimentally resolved [3].

In evolutionary biology, fitness landscapes have been used to study adaptation. In a strong selection weak mutation regime [6], the evolutionary paths followed by the populations are restricted to paths of increasing fitness. In this perspective, it has been emphasized that many fundamental features of adaptation depend on whether the landscape is smooth or rugged. Among other things, the ruggedness and the properties of fitness landscapes have been related to speciation processes [7, 8], to the benefits of sexual reproduction [9, 10, 11, 12], and more generally, to the repeatability of the adaptation process (e.g. [13, 14, 15, 16]).

One of the most basic ingredients that characterize the structure of fitness landscapes is epistasis. Epistasis can be broadly defined as the interaction between the effects of mutations at different loci. It is usually defined as the non-multiplicative part of the fitness effects of combined mutations, that is the non-additive part, in log-scale. In the presence of epistasis, the fitness effect of a mutation at a given locus depends on the genetic background and consequenty, a mutation at a given locus changes the distribution of fitness effects of other mutations at other loci. For the 2-loci 2-alleles case, assuming that the genotype with the smallest fitness is labeled 00, epistasis can be expressed (in logscale) as the departure from additivity: e = *ƒ*(11) − *ƒ*(10) − *ƒ*(01) + *ƒ*(00), where *ƒ*(*ij*) is the malthusian fitness of the genotype ij (e.g. [17]).

Assuming random fitness values (*i.e*. NK landscapes), Kauffman [13] showed a positive correlation between the amount of epistatic interactions and the ruggedness of a landscape, defined as the density of peaks (genotypes with no fitter neighbors). As the number of loci that interact together grows, the landscape is more rugged, and more local peaks end evolutionary paths. At the maximum number of epistatic interactions, the fitnesses of each genotype are completely uncorrelated, resulting in a so-called House-of-Cards model [18, 19], for which several measures on paths of increasing fitness were derived [19, 20, 21, 22].

Other measures such as *r/s* ratio (ratio of the roughness over additive fitness), fraction of sign epistasis, fraction of nonlinear interactions in the Fourier spectrum of the landscape, number of accessible paths to the maximum (see detailed description in the Appendix) were also proposed to characterize the structure of fitness landscapes. As they all represent direct or indirect measures of epistasis, all these measures were shown to be pairwise correlated in experimental fitness landscapes [23].

Quite surprisingly, all these global measures of epistasis are only indirectly related to the simplest definition of epistasis, namely the interaction of other mutations with the fitness effect of a specific mutation. The measure that is most related to this definition is the fraction of sign and reciprocal sign epistasis, but this measure is sensitive only to strong epistasis, *i.e*. mutations changing from beneficial to deleterious in different backgrounds. Other finer measures like *r/s* and the fraction of nonlinear interactions in the Fourier spectrum are actually more related to the global deviations from linearity and their contributions to fitness variance than to the definition above. Therefore, it is worth exploring the possibility of finding a natural measure of epistasis that follows the definition above more closely.

Here, we describe new measures that can be used to characterize epistasis and structure of fitness landscapes (Figure 1). In section 2.1 we present our basic measure of the amount of epistasis, γ, that is the single-step correlation of fitness effects for mutations between neighbor genotypes (Figure 1b). It is a direct measure of epistasis, *i.e*. it measures how much the fitness effect of a mutation is affected when a genotype experiences another mutation. We also discuss how it can be applied to landscapes with missing data. In section 2.3 we discuss its properties. As all correlation measures, γ ranges from –1 to +1 and is a very natural quantity to describe the amount of epistasis in the landscape. In section 2.4 we show how to apply the measure to individual mutations and in section 2.5 we extend it to landscapes where only the signs of fitness effects of mutations are known.

In section 2.6 we discuss a larger class of measures, *i.e*. the correlation of fitness effects of mutations across multiple mutations, and we study its dependence on mutational distance. This dependence contains information on the epistatic effects of multiple mutations and depends both on the amount and on the nature of epistatic interactions. In section 2.7 we compute analytically the mean of these quantities for a variety of landscape models (the House of Cards, the Rough Mount Fuji and NK landscapes as well as Ising and Eggbox models).

Finally, in section 3 we discuss the relation of γ with previous measure of epistasis and in section 4 we show how the new measures can be used to understand better the features of experimental landscapes.

**Figure 1:**
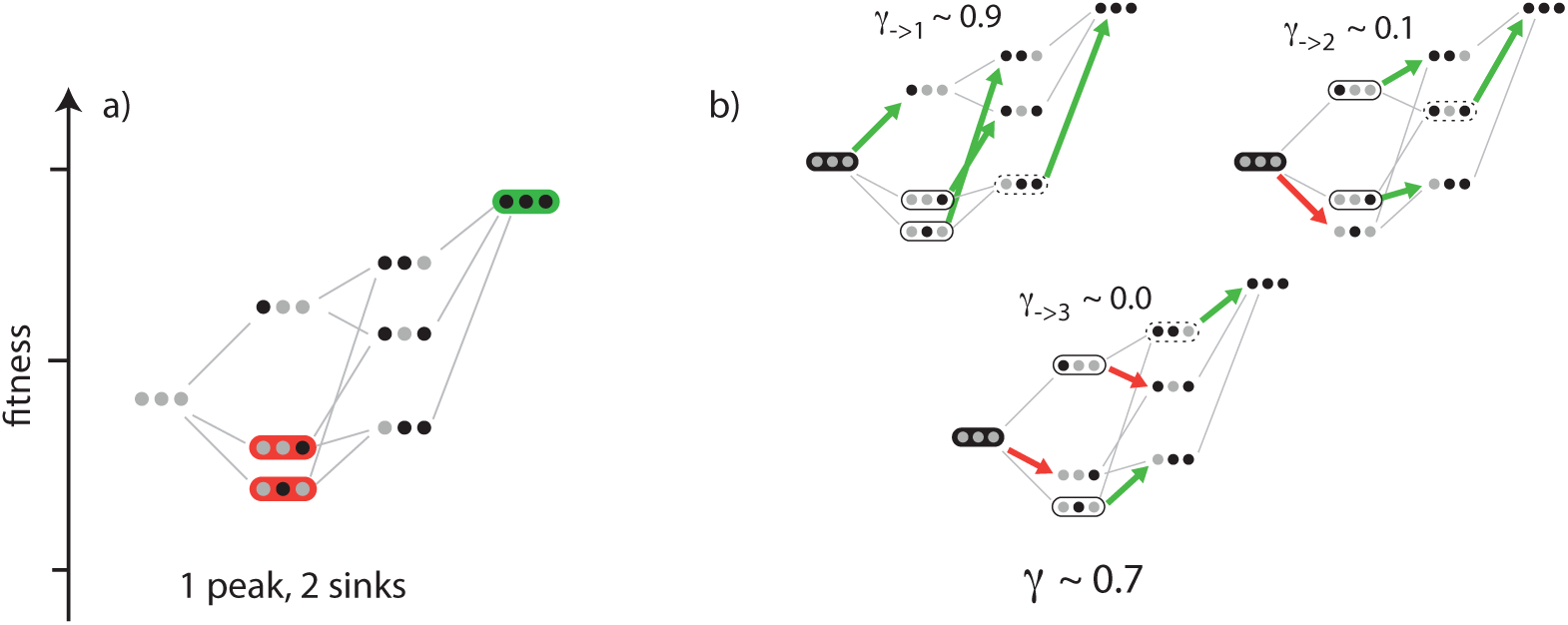
Measures of epistasis in fitness landscapes. We depict two measures on the same fitness landscape (of 3 loci with 2 alleles each). (a) Peaks, here in green, are genotypes with no fitter neighbors (whereas sinks are genotypes with only fitter neighbors). The number of peaks is a classical measure of epistasis. (b) γ is the pairwise correlation in fitness effect of mutation between neighbor genotypes. It measures how much another mutation in a genotype affects the focal mutation, averaged across all mutations and the whole landscape. Here the average correlation is good (γ ≈ 0.7). γ_→*i*_ is the correlation in fitness effect of mutation *i* between neighboring genotypes. In the example, mutations at locus 1 are almost independent of the genotypes (γ_→1_ ≈ 0.9), whereas the effects of the mutations at locus 3 show almost no correlation across genotypes (γ_→3_ ≈ 0).

## 2. Epistasis as correlation of fitness effects: γ

### 2.1. Definition

In this section, we will derive and discuss a new measure that is a natural description of the amount of epistasis in fitness landscapes. This new measure, denoted by γ, is simply the correlation of fitness effects of the same mutation in single-mutant neighbors (see Figure 1b and Figure 2). It measures how the effect of a focal mutation is altered by another mutation at another locus in the background, averaged across the whole landscape.

**Figure 2:**
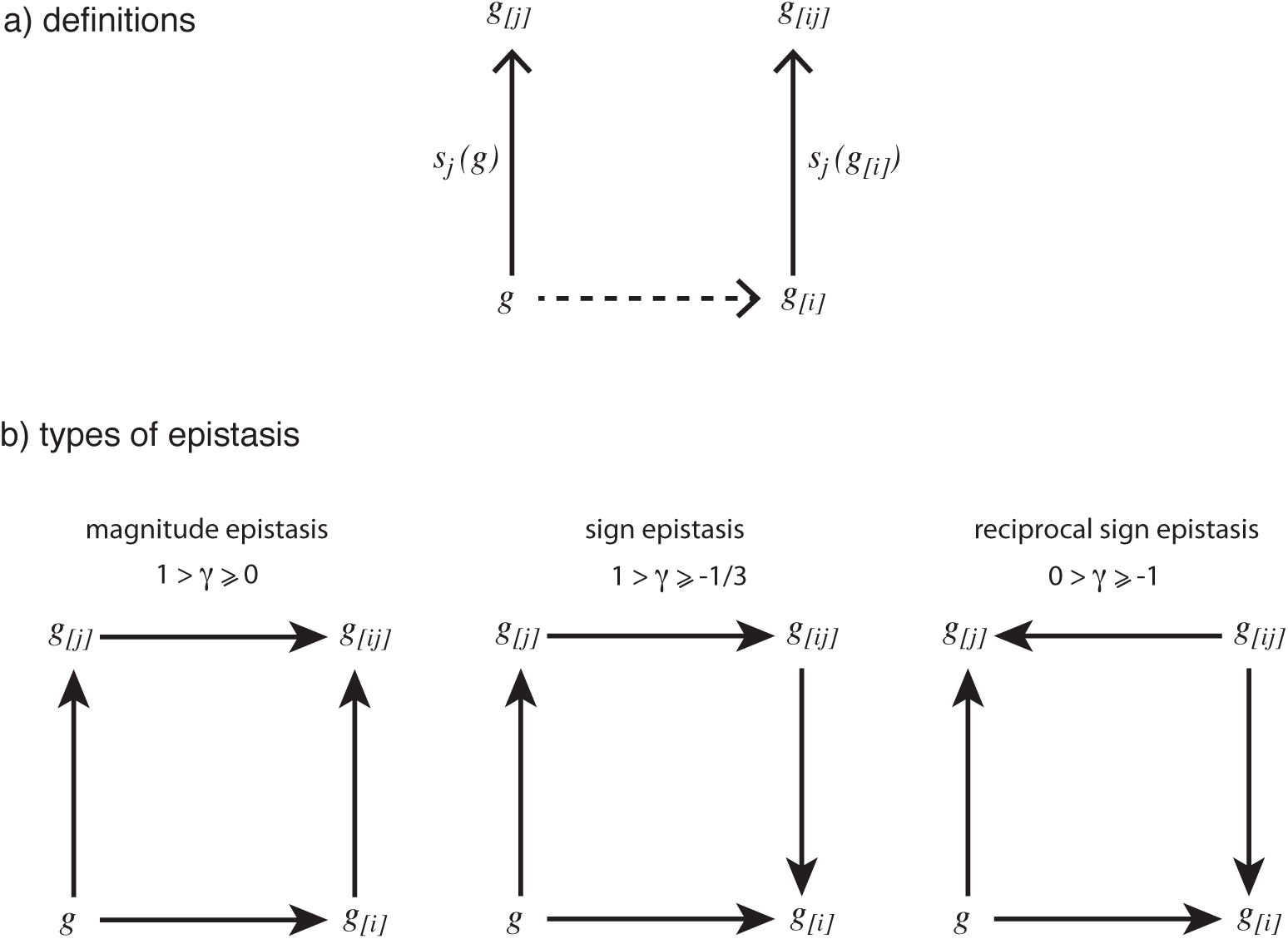
(a) Notation: γ is the correlation between the fitness effects *s*_*j*_(*g*) and *s*_*j*_(*g*_[*i*]_) over all genotypes *g* and mutations *i, j* in the landscape. (b) Types of epistasis, possible values of γ and examples of the corresponding fitness graphs. In Figure (b), fitness increases in the direction of the arrows.

In the following, we will define it properly in mathematical terms for the bi-allelic case. We denote the (log-scaled) fitness of a genotype *g* by *ƒ*(*g*). We also define *g*_[*i*]_, the genotype *g* where the locus *i* is mutated. The fitness effect of a mutation at locus *j*, *i.e*. the log-scale selection coefficient of the mutation, is denoted by *s*_*j*_(*g*) = *ƒ*(*g*_[*j*]_) – *ƒ*(*g*). The new measure γ is then defined as the correlation between two fitness differences *sj(g)* and *s*_*j*_(*g*_[*i*]_) measured from genotypes that are one mutation away, as illustrated by Figure 2a.

Noting that the average of *s*_*j*_(*g*) across all genotypes and mutations in the landscape is 0, we define γ as:

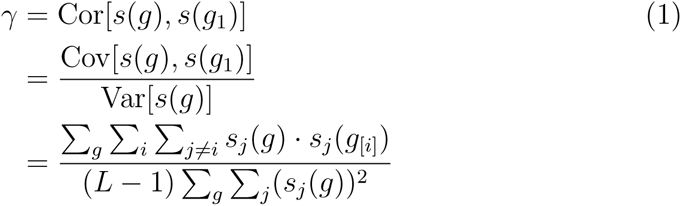

where *g*_1_ indicates a generic genotype that differs from *g* by a single mutation. For multiallelic landscapes, the same definition γ = Cor[s(*g*), *s*(*g*_1_)] can be immediately generalized to any number of alleles.

### 2.2. Landscapes with missing data

Even though γ is originally defined in term of fitness effects of the mutations, it can be easily recomputed by only using the fitness values themselves. If we denote *ρ*_d_ = Cor[*ƒ*(*g*), *ƒ*(*g*_*d*_)] the fitness correlation function at Hamming distance *d*, i.e. the correlation between the fitness of genotypes *d* mutations apart, it is possible to rewrite γ by the simple formula (see proof in the Appendix):

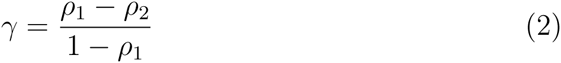

Besides its general interest, this formulation allows us to measure γ in the presence of missing data. Indeed, fitness correlation functions do not need fitness data for all combinations of some set of mutations in order to be estimated.

The above formula can be applied in a straightforward way to landscapes with missing data, by considering only pairs of genotypes with known fitness in the computation of the correlations. The result provides an estimate of the value of γ for the full landscape. However, the error of the estimate increases with the fraction of genotypes of unknown fitness. For sparse landscapes, an estimate can be obtained only if the fitness is known for a sufficient number of pairs of genotypes at distance *d* = 1 (for *ρ*_1_) and 2 (for *ρ*_2_). Moreover, these pairs should be randomly distributed across the landscape.

### 2.3. Interpretation

To make clear that the above measure is a metric of epistasis, we rewrite the above equation as

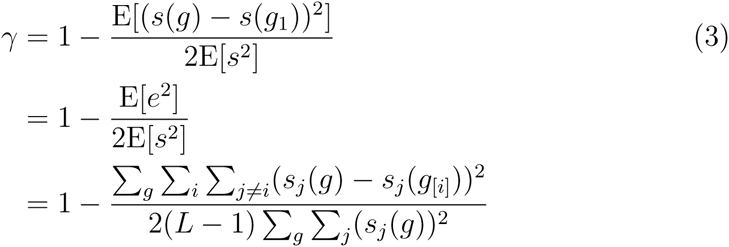

where *e* is the usual measure of epistasis defined in the Introduction and E[*x*] denotes the expected value of the quantity *x* over genotypes and mutations.

When there is no epistasis, the fitness effects do not depend on the background and γ = 1, *i.e*. perfect correlation between fitness effects. The deviation of γ from 1 is proportional to the square of *e*_*ij*_ = *ƒ*(*g*_[*ij*]_) – *ƒ*(*g*_[*i*]_) – *ƒ*(*g*_[*j*]_) + *ƒ*(*g*), which is a standard measure of the amount of epistatic effect (for 2-alleles 2-loci), normalized by the average squared fitness effect. Thanks to its normalization, this measure of epistasis is not affected by the scale and the absolute level of fitness, but only by relative differences in fitness. Shifting fitnesses by a multiplicative or additive factor does not change this measure.

The measure γ is defined as a correlation, therefore it is bounded by – 1 ≤ γ ≤ 1, with γ = 1 in the case of no epistasis. The value of γ is related to the prevalent type of epistatic interactions (see proof in the Appendix), which are summarized in Figure 2b:

- magnitude epistasis refers to pairwise interactions that do not change the signs of fitness effects. Magnitude epistasis would still result in a positive correlation between fitness effects, therefore γ would still be positive even if smaller than 1: 1 > γ ≥ 0;
- sign epistasis refers to pairwise interactions where the fitness effects of one mutation change sign after the other mutation. Sign epistasis would contribute with terms of both signs to the correlation, therefore resulting in values centered around 0: 1 > γ ≥ –1/3;
- finally, reciprocal sign epistasis refers to pairwise interactions where both fitness effects change sign. Reciprocal sign epistasis would imply a negative correlation between fitness effects, and therefore a negative value of γ: 0 > γ ≥ – 1.

The deviation of the mean value of γ from 1 for simple landscape models measures epistasis as a function of the parameters of the models (see section 2.7 for more details). For example, in NK landscapes, the epistasis grows with the parameter K describing the number of loci involved in each interaction and, in fact we have the approximate equation

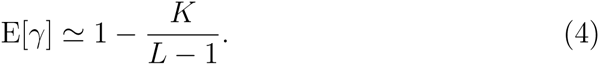

For the House of Cards (HoC) model, *i.e*. a maximally uncorrelated landscape, we have *K* = *L* – 1 and therefore

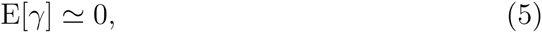

*i.e*. this model shows strong random epistasis.

For Rough Mount Fuji (RMF) models, which are combinations of an additive landscape and a completely uncorrelated one, the correlation of fitness effects is

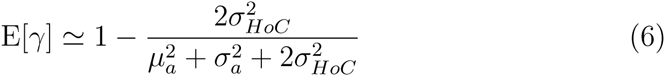

where *μ*_*a*_ and σ_*a*_ are the mean and variance of the additive fitness effects and 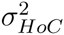 is the variance of the uncorrelated HoC component. Therefore, in this case, the measure of epistasis is proportional to the variance contribution of the uncorrelated component.

### 2.4. Epistasis for specific mutations

The correlation of fitness effects is also a useful measure of the interaction between specific mutations. Some simple generalizations of the γ measure are:

- γ_*i*→_, which describes the epistatic effect of a mutation in locus *i* on other loci:

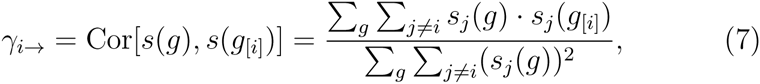
- γ_→*j*_, which describes the epistatic effects of other mutations on locus *j*:

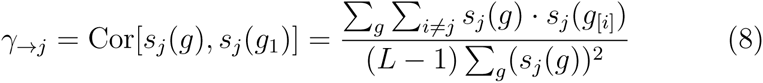
- γ_*i*→*j*_, which is a matrix that describes the epistatic effect of locus *i* on locus *j*:

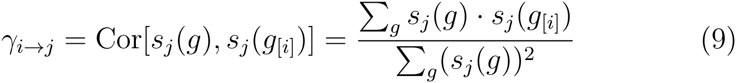

These measures can also be generalized easily to multiallelic landscapes by considering pairs of mutations at different loci.

The measure γ_*i*→_, γ_→*j*_ and especially γ_*i*→*j*_ are useful for exploratory and illustrative purposes, since they summarize the interactions between mutations in a clear and compact way, as it can be seen in Figure 4.

**Figure 4:**
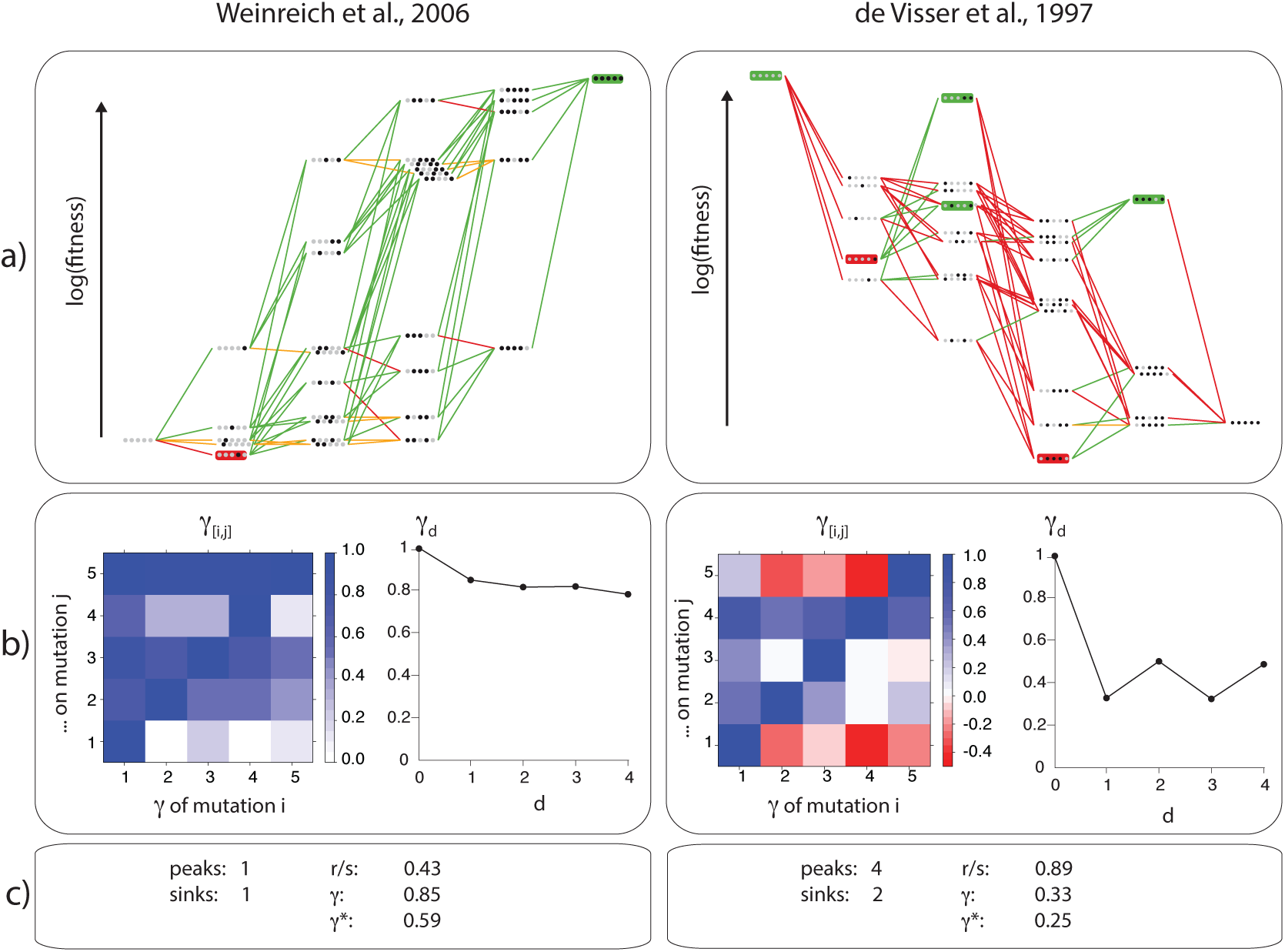
Values of several measures applied to two experimental landscapes. a) Illustrations of the landscapes using Magellan [25]. b) (left) Interactions between pairs of mutations γ_*i*→*j*_: blue = no interaction, white = strong random interaction, red = strong interaction in sign; (right) Decay of γ_*d*_ with Hamming distance. c) Measures for the landscape.

It is also possible to use the more direct measure 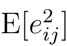 as an alternative to γ_*i*→*j*_. The difference lies in the normalization: 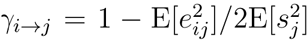, therefore γ_*i*→*j*_ treats both large and small mutations in the same way while 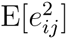 is larger for large mutations. The choice of the most appropriate measure depends on the question, *i.e*. if the focus is on the interactions across all mutations, or only the largest ones.

### 2.5. Correlation in signs (γ*)

In many experimental situations, fitness is not clearly measurable on an absolute scale, but it is possible to rank the genotypes in order of increasing fitness, or at least to state if a mutation is deleterious or beneficial.

The fitness landscape can be then represented as an acyclic oriented graph, *i.e*. an oriented network where links between genotypes represent single, fitness-increasing mutations. Hereafter, we will refer to this graph as the fitness graph. As an example, the fitness graphs corresponding to different types of epistasis for 2 loci are illustrated in Figure 2b.

In this context, it is still possible to measure epistasis via the same method by employing a modified measure γ* which uses just the sign of the fitness effects, instead of their value. We define 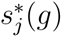 as the sign of *s*_*j*_(*g*). A more robust variant would be

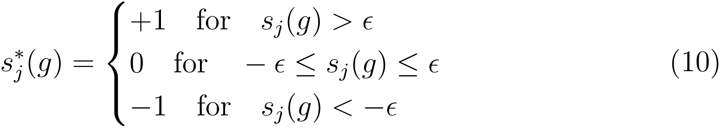

where *ϵ* is a tolerance parameter (possibly depending on the genotype, and larger than the experimental errors). The measure γ* is defined as before:

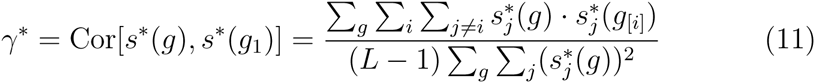

If the landscape has no neutral mutations, we can show that this measure is related to other commonly employed measures for fitness graphs. Consider all possible pairwise mutational motifs in the fitness graph and classify the type of epistasis in each motif as magnitude epistasis, sign epistasis and reciprocal sign epistasis (see Figure 2b). We denote the fraction of motifs in each class by *ϕ*_*m*_, *ϕ*_*s*_ and *ϕ*_*rs*_ respectively. We have the relation (see proof in the Appendix)

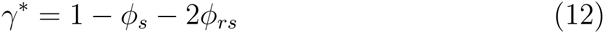

What is even more interesting is that both in models and in real landscapes, the results of γ and γ* are often numerically close and highly correlated (see below). The only exception is represented by landscapes with weak epistatic interactions dominated by magnitude epistasis, where γ* = 1. This suggests that γ* could be used in place of γ for landscapes where only fitness ranks are known. These measures represent therefore a bridge between fitness graph-based measures and quantitative measures based on absolute fitness.

### 2.6. Decay of the correlation with distance

The γ measure provides information on the amount of epistasis but cannot discriminate between different types or models of fitness landscapes, as it occurs for any single measure of the amount/strength of epistatic interactions. In fact, there are many landscapes with widely different structure but with the same γ. For example, a HoC model realization would have γ = 0 as would a landscape composed by an equal mixture of additive and reciprocal sign epistatic interactions (like in an EMF model).

However, a natural and interesting extension of this measure is given by the full decay of the correlation of fitness effects with distance *d*, which correspond to the cumulative epistatic effect of *d* mutations and can be defined as:

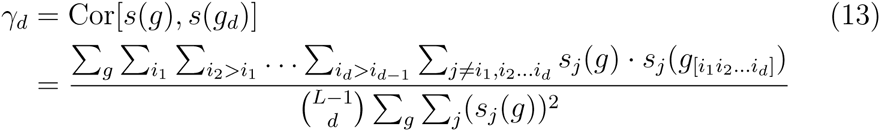

where γ_1_ = γ. As with γ, γ_d_ can be expressed in terms of the fitness correlation functions at distance *d*, *ρ*_*d*_:

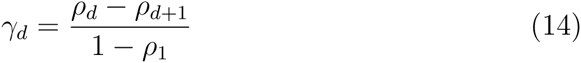

The decay of γ_*d*_ with the Hamming distance *d* is an interesting object of study in itself, since it describes how the epistatic effects of different mutations interact with each other and their cumulative effect. The mean of γ_*d*_ can be computed analytically in most models of fitness landscapes and it brings extra information on the structure of the landscape. Different models have a different behaviour (Figure 3): RMF and HoC models show an abrupt fall already at *d* = 1 and then a flat profile, while NK models have a gradual, approximately exponential decay with rate *K*/(*L* – 1) (Supplementary Figure 1). The Ising model (based on pairwise reciprocal sign epistasis) decays linearly until –1, while the eggbox (maximally epistatic, anticorrelated) oscillates between –1 and 1.

**Figure 3:**
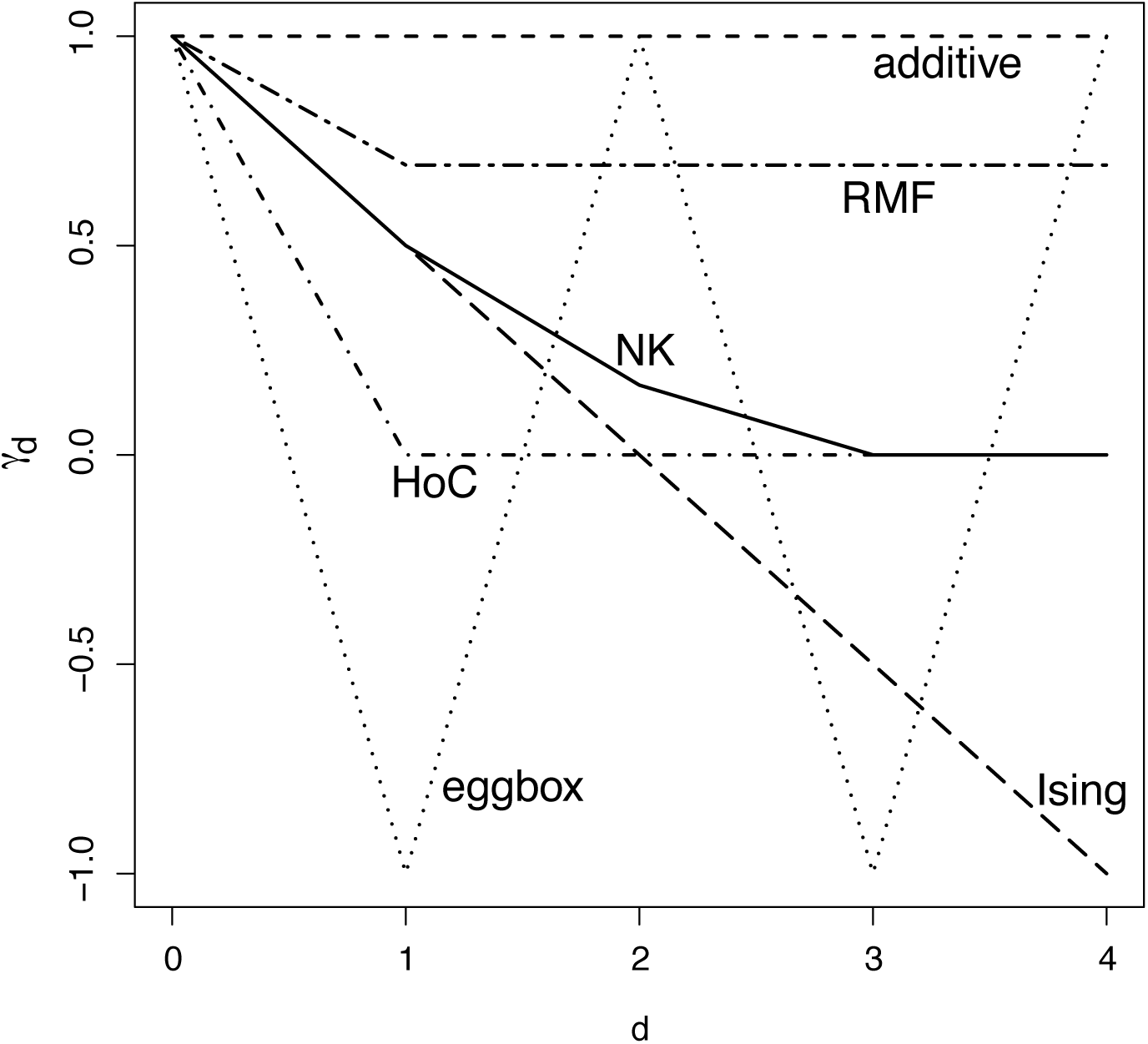
Behaviour of the average correlation of fitness effects γ_*d*_ at different distances in model landscapes with *L* = 5. The NK landscape has *K* = 2 and the RMF is a mixture of 60% additive component and 40% HoC.

### 2.7. Formulae for γ_*d*_ in model landscapes

Notation: a genotype *g* is a biallelic sequence of length *L* of alleles *g* = (*A*_1_*A*_2_*A*_3_…*A*_*L*_) with *A*_*i*_ ∈ {0,1}. We assume an observed fitness *ƒ*(*g*) given by the model

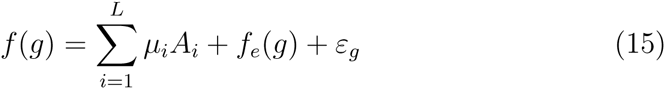

that is, an additive contribution 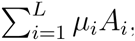, an epistatic contribution *ƒ*_*e*_(*g*) from some fitness model, plus the effect of measurement errors *ε*_*g*_. We assume these errors to be unbiased and uncorrelated: E[*ε*_*g*_] = 0, 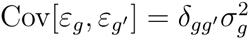 where δ_*gg*′_ is the Kronecker delta, *i.e*. δ_*gg*′_ = 1 when *g* = *g*′ and 0 otherwise.

We define the mean squared additive effect 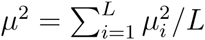 and the mean squared experimental error 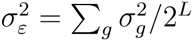.

The expected value of γ_*d*_ can be computed approximately by taking the ratio of the expected values of numerator and denominator of eq. (13) rearranged as eq. (3), instead of the expected value of the ratio (the ≃ sign in all our formulae refers to this approximation). The result is

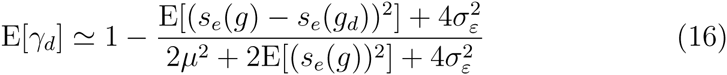

with *s*_*e*_(*g*) = *ƒ*_*e*_(*g*_[*j*]_) − *ƒ*_*e*_(*g*) being the analogue of *s*(*g*) restricted to the epistatic contribution. Note that E[(*s*_*e*_(*g*)–*s*_*e*_(*g*_*d*_))^2^] = 2(1–E[γ_*e*_*d*__])E[(*s*_*e*_(*g*))^2^] if γ_*e*_*d*__ is the γ_*d*_ statistic of the epistatic contribution *ƒ*_*e*_(*g*).

#### 2.7.1. Additive model

In these models *ƒ*_*e*_ = 0 and the only reduction in correlation is due to experimental noise:

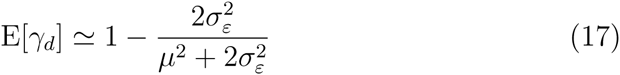

#### 2.7.2. RMF and HoC models

In these models, *ƒ*_*e*_(*g*) corresponds to the HoC model, *i.e*. they are i.i.d. random variables. Denote by 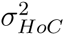 the variance of the distribution of fitnesses in the HoC model:

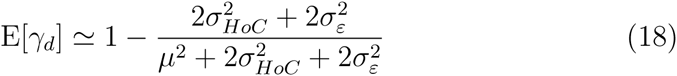

Since 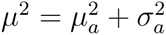 for a Gaussian distribution of additive fitness effects, we obtain equation (6) for *d* = 1 and 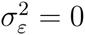.

#### 2.7.3. NK models

In these models, 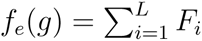 where the *F*_*i*_s are i.i.d. random variables that depend on *i* and other *K* indices, randomly chosen. The *F*_*i*_s have mean *ƒ*_0_ and variance 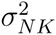.

The fitness correlation function *ρ*_*d*_ is known exactly for the pure NK model [24]: 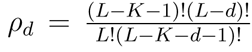 With eq. (14) it is straightforward to compute E[γ_*e*_*d*__] from this. The variance E[*s*_*e*_(*g*)^2^] is given by 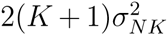 because on average *K* + 1 of the *F*_*i*_ change in a single mutation, each of these differences having twice the variance of the fitness contribution (since the variance of the difference of two i.i.d. variables is twice their variance). The final result is

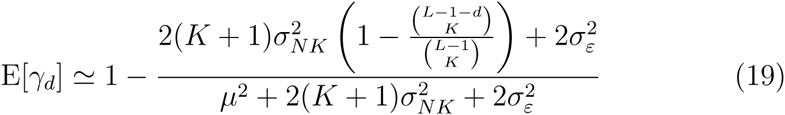

Substituting 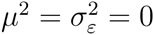 and *d* = 1 yields equation (4).

#### 2.7.4. Ising model

In the following, we define *S*_*i*_ = 2*A*_*i*_ –1 ∈ {–1, +1}. Using the above notations, in these models, *ƒ*_*e*_(*g*) = ∑_*i*_ *J*_*i*_*S*_*i*_*S*_*i+1*_ where the incompatibility coefficients *J*_*i*_ are randomly extracted from a Gaussian distribution with mean *μ*_*c*_ and variance 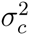. We define 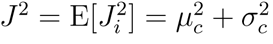.

A mutation at locus *i* will invert contributions of the terms containing *J*_*i*_ and *J*_*i*-1_ adding 8*J*^2^ to E[(*s*_*e*_(*g*))^2^]. At the edge of the genome, loci interacts with only one neighbor, so this is reduced by a factor of 2, thus 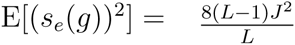.

Only mutations at *j* – 1 and *j* + 1 affect the value of *s*_*j*_(*g*) – *s*_*j*_(*g*_*d*_). Choosing *d* mutations out of *L* – 1 and applying the hypergeometrical distribution, there are probabilities 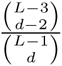 and 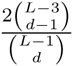 to choose both or exactly one of them, respectively. Each relevant mutation changes the effect of mutating *j* by ±4*J*. Thus (ignoring boundaries) 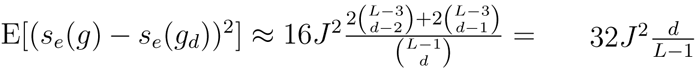. On the boundary only one mutation can influence *s*_*j*_, resulting in a reduced contribution of 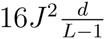 and so together:

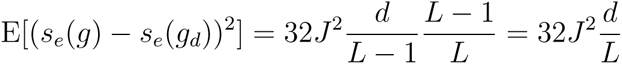

and therefore the result is

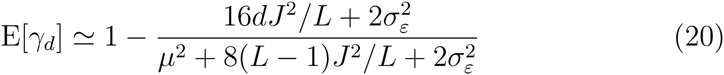

More general results can be obtained for a generic spin glass model 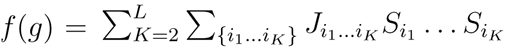 where {*i*_1_… *i*_*K*_} are ordered sets. We define 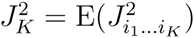 and 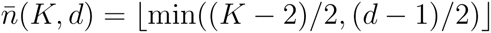.

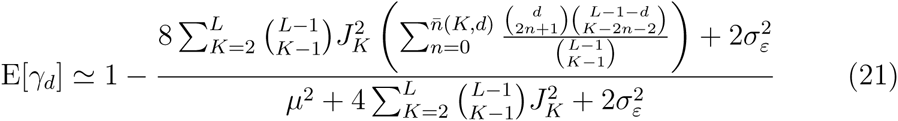

#### 2.7.5. Eggbox model

In this model, 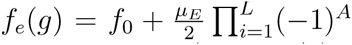, i.e. each mutation switches the fitness from the highest value (*ƒ*_0_ + *μ*_*E*_/2) to the lowest (*ƒ*_0_ – *μ*_*E*_/2) or the other way. The difference in the epistatic fitness effects from two genotypes separated by an odd number of mutations is ±*μ*_*E*_, while it is 0 from two genotypes separated by an even number of mutations, therefore

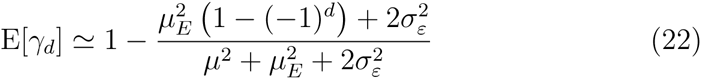

#### 2.7.6. Block models

We define a general block model with *B* blocks by the fitness 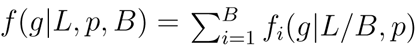 where each contribution *ƒ*_*i*_(*g*) comes from an i.i.d. block of length *L/B* described by any model (NK, Rough Mt. Fuji, etc) with parameters *p*. Denote by 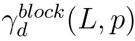 the average correlation of fitness differences for the pure model. The average correlation of fitness differences for the full block model can be obtained as

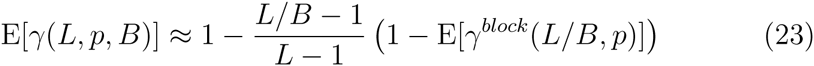

so the block structure reduces the epistasis roughly speaking by a factor 1*/B*, which corresponds to the probability that the second mutations act on the same block as the first.

Similarly, the general result for the decay with distance is

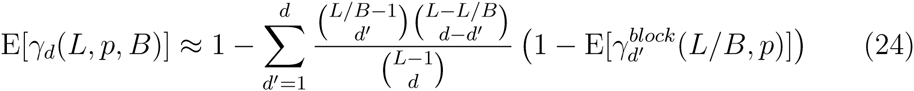

where the hypergeometric distribution counts the number of mutations in the same block of the first mutation.

## 3. Relations between measures of epistasis

In this section we discuss the relations existing between the newly proposed measures of epistasis and the existing ones.

We expect that γ and γ* would be correlated to other measures of epistasis. In fact, we already discussed how they are related to some of the existing measures. In particular, (i) for pairs of loci, γ is directly related to the common definition of epistasis e as 1 – γ ∝ *e*^2^: in fact, for the whole landscape we have γ=1 – E[*e*^2^]/2E[*s*^2^], while for a pair of mutations 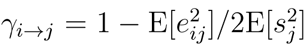; (ii) for the whole landscape, γ can be rewritten as a function of the fitness correlation functions *ρ*_*d*_ as γ_*d*_ = (*ρ*_*d*_ – *ρ*_*d*+1_)/(1 – *ρ*1); (iii) γ* is directly related to the number of square motifs with sign and reciprocal sign epistasis as γ* = 1 – *ϕ*_*s*_ – 2*ϕ*_*rs*_.

Furthermore, it is possible to show that γ is a function of the Fourier spectrum of the landscape, provided that the standard orthonormal basis is used for the Fourier series. If we denote by *W*_*J*_ the normalized weight of the coefficients of order *J* in the Fourier spectrum, *i.e*. the sum of the squared coefficients of all *J*-loci interactions normalized by the sum of all squared coefficients, the relation is

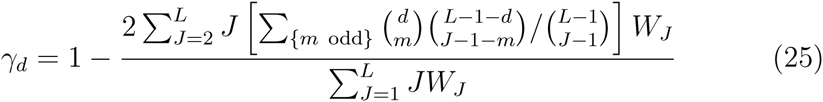

(see proof in the Appendix). Our measure of epistasis is therefore

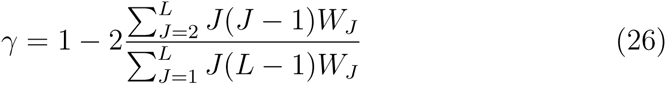

which resembles the usual measure of epistasis from Fourier expansion, *i.e*. the fraction of non-linear interactions in the Fourier spectrum 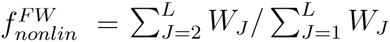 [23], showing again the close relation with previous measures of epistasis. The main difference is the weight of higher-order interactions: the contribution of J-loci interactions to γ grows like *J*^2^ for large *J*, so that the effect of interactions is stronger if they involve more loci.

To evaluate in a more systematic way the relations between these and other measures, we perform a correlation analysis similar to [23] but using models instead of experimental landscapes. We select the number of peaks, the roughness/slope ratio (ratio between epistatic “noise” and additive component, see Appendix), γ and γ* as measures of epistasis. We compute the Spearman correlation coefficients of all pairs of measures in the RMF landscape model varying the model parameters (in particular, the ruggedness).

The pairwise correlations (Table 1) confirm the intuition that the measures related to epistasis are all strongly correlated, the strongest correlation being between γ and γ* as expected.

Similar conclusions apply to the Ising and eggbox models with a “Mount Fuji” component (IMF and EMF), which represent different kinds of epistatic interactions. Interestingly, γ seems to be strongly related to *r/s* in these models while γ* is highly correlated with the number of peaks and sinks. This is probably due to the extreme compensatory interactions present in these models.

**Table 1:**
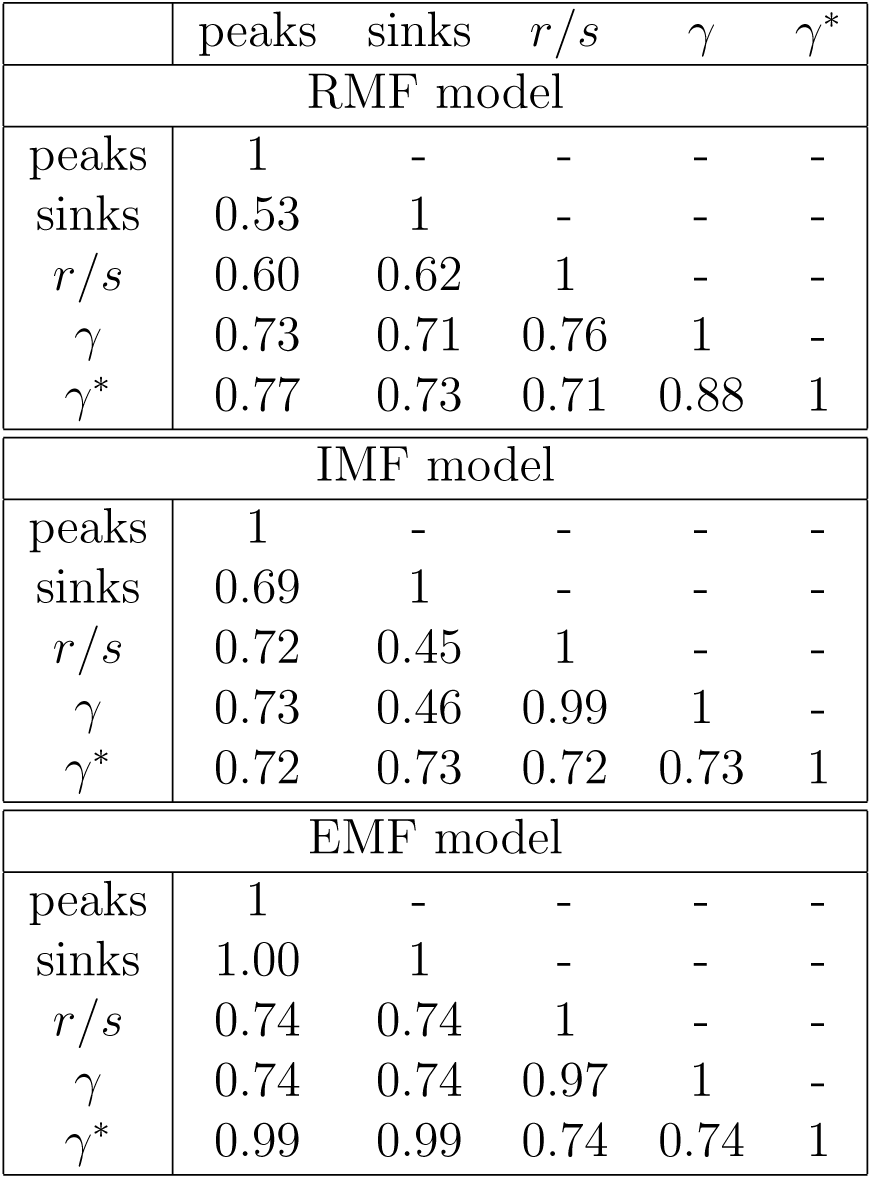
Spearman *ρ*^2^ correlation of pairs of measures across 10^4^ realizations of the RMF, IMF and EMF models with *L* = 5, σ>_*HoC*_ = 1 (for RMF), *μ*_*I*_ = 1 and σ_*I*_ = 0.2 (for IMF), *μ*_*E*_ = 1 and σ_*E*_ = 0.2 (for EMF), σ_*a*_ = *μ*_*a*_/10 and *μ*_*a*_ log-uniformly distributed in [0.01,10].

Each measure of epistasis has its own advantages and disadvantages. γ has several interesting features:

- γ has a natural and direct interpretation in terms of epistasis. Other measures (number of peaks and number of accessible paths) are only indirectly related to epistasis and more focused on evolution on the landscape.
- γ is a fine-grained and robust measure of epistasis. It is less noisy than discrete measures like the number of peaks, and it is able to measure also magnitude epistasis unlike fitness graph-based measures like *ϕ*_*s*_, *ϕ*_*rs*_.
- γ weights epistatic interactions according to their structure, not only their strength. A complex interaction involving many loci in a landscape is weighted much more than a single compensatory interaction between two loci, even if their fitness variance is the same. This does not occur with measures like *r/s* or the fraction of non-linear interactions 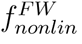 in the Fourier-Walsh spectrum. In fact, as discussed above, γ weights the spectrum according to the square of the number of loci involved in the interaction. On the other hand, the usual spectrum measure 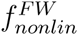 weights all interactions equally.

The last point is probably the most important reason to use γ in addition to other statistics. In practice, *r/s*, 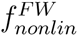 are mostly sensitive to the global epistatic contribution to the variance, so even a single pairwise epistatic interaction of large effect would result in a strong signal. In contrast, γ would show strong epistasis only if the interaction would involve many loci, while for a pairwise interaction it would show weak epistasis even if the interaction would dominate the fitness variance.

As an example, a landscape of size *L* with additive fitness effects *s*^2^ and a single compensatory interaction of strength e between locus 1 and 2 has 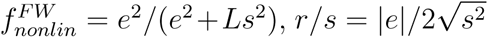 and γ=1 – 16*e*^2^/[(8*e*^2^ + *Ls*^2^)(*L* – 1)]. For example, for *L* = 5 and *s*^2^ = *e*^2^, we have 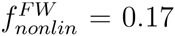, *r/s* = 0.5 and γ = 0.7. The same landscape, but with a maximal compensatory interaction (eggbox) of strength e across all loci, has the same values of 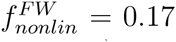 and *r/s* = 0.5 but a negative correlation (*i.e*. much stronger epistasis) γ = 1 – 8*e*^2^/(4*e*^2^ + *s*^2^) = –0.6.

## 4. Epistasis in two experimental landscapes

As an example of application of the new measures, we use them to analyze two complete experimental landscapes of size *L* = 5. These landscapes are illustrated in Figure 4.

The first landscape is the landscape of antibiotic (cefotaxim) resistance of β-lactamase mutations in an *Escherichia coli* plasmid from Weinreich et al. [26] (Figure 4 left). The 5 mutations have a very strong effect that together give a 4 × 10^4^ increase in antibiotic resistance and were therefore selected together. Given the huge selective advantage of the combined mutations, this landscape is single-peaked, where the peak corresponds to the five-point mutant. It also has a single sink, that interestingly does not correspond to the wild type.

The second is one of the four *L* = 5 complete sublandscape (csI) [20] of a larger landscape (*L* = 8) of deleterious mutations in *Aspergillus niger* from de Visser et al. [27] (Figure 4 right). This landscape is a combination of unrelated deleterious mutations where epistatic interactions were not filtered by natural selection. This landscape has 4 peaks and 2 sinks; in fact, at present it is one of the most rugged among the completely resolved landscapes.

As the landscapes were derived in completely different settings (co-selected beneficial for β-lactamase and random deleterious for *Aspergillus*), we might not be surprised to find that these landscapes exhibit very different structures. Indeed theoretical arguments support the intuition that landscapes of co-selected mutations differ radically from landscape of random mutation [28, 29, 30]. The difference in ruggedness between β-lactamase and *Aspergillus* landscapes is confirmed by the values of γ (0.85 *vs* 0.33) and *r/s* (0.43 *vs* 0.89).

To further explore the landscapes, we compute the γ_*i*→*j*_ matrices to illustrate and summarize the interactions between mutations (Figure 4b). In the β-lactamase landscape, there are some clear interactions between mutations (between the 2nd or the 4th and the 1st mutations, or between the 5th and the 4th) but none of these interactions is characterized by strong sign epistasis (no red cell). On the other hand, the *Aspergillus* landscape contains several examples of interactions dominated by strong sign epistasis (for example, between the 2nd or the 4th and the 1st or the 5th mutations).

Similar conclusions come from the analysis of the decay of γ with distance. The decay in the *β*-lactamase landscape is immediate but decays slowly after the first mutation (Figure 4b left), resembling the behavior of RMF models.

An interesting example of the power of γ_*d*_ is represented by the *Aspergillus* landscape (Figure 4b right). This landscape shows a non-monotonic decay, with correlation γ_*d*_ bouncing up and down. This points to a compensatory structure of reciprocal sign epistasis, which is not only due to pairwise compensation, but extends to distance 4, *i.e*. to the whole landscape. In fact, the behavior of γ_*d*_ suggests a mixture of an RMF landscape and an extreme case of compensatory interactions, like the Eggbox model. This surprising result does not come out in a straightforward way while looking at other measures, even when looking at the Fourier spectrum [31]. Indeed, although the coefficient of the highest order in the Fourier decomposition measures the amount of eggbox, it compares to coefficients of smaller orders that have a complex intermingling when epistasis is not purely reciprocally signed at all orders. These coefficients contribute as well to the behaviour of γ_*d*_.

## 5. Discussion

In this work, we presented a new set of landscape measures of epistasis which have a simple interpretation and cover a range of potential applications. These measures among others have been implemented in MAGELLAN, a graphical tool to explore small fitness landscapes [25].

The first application is the measure of epistasis in a comparable way across landscapes. The correlation of fitness effects γ is a natural measure for this. This measure can be used also for pairs of mutations, to explore the strength of epistatic interactions between mutations in a compact way. While γ is a natural and direct measure of epistasis, there are other possible measures, and different measures have different strengths. The interpretation of γ is clear: as correlation of fitness effects, it measures how epistatic interactions affect the fitness effect of a single mutation. The alternative measures of epistasis *r/s* and 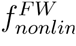 should be used instead when the focus is on the epistatic contribution to fitness variance in the landscape. Loosely speaking, γ measures local epistasis while *r/s* and 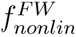 measure global epistasis.

In terms of γ, there is a natural scale for the strength of epistatic interactions, from purely additive interactions (γ = 1), through random interactions (γ = 0, e.g. HoC) to a fully compensatory landscape (γ = – 1, e.g. eggbox). Interactions in landscapes with γ < 0 are dominated by strong sign and reciprocal sign epistasis between most loci, therefore we expect such landscapes to be rare and possible only for some sets of mutations, as selection tends to favor mutations with positive interactions. In fact, the two experimental fitness landscapes analyzed have positive values of γ. Yet, the amount/strength of epistasis in the landscape by de Visser et al. is remarkably high: γ = 0.33 means that the fitness effect of a mutation in a given genotype, is a poor predictor of the fitness effect of the same mutation in a neighbor genotype that only differs by a *single* mutation.

The quantity γ can only be defined for genotype spaces where for each mutation from a given genotype, there is a corresponding mutation from each of the neighbour genotypes (except the final genotype of the mutation itself). This is the case of nucleotide or protein sequences, which underlie most molecular fitness landscapes. Note that the presence of a mutation in the landscape has nothing to do with its fitness effect being measured or not; in fact, γ is well defined also in DNA or protein landscapes with missing data, even if in this case the fitness data allow only an uncertain estimation of it.

From the mathematical point of view, there is a natural geometric interpretation for γ in terms of differential geometry on lattices. In fact, γ reinterprets the epistasis as e_*ij*_ = *s*_*j*_ (*g*_[*i*]_) – *s*_*j*_(*g*), that is the local parallel transport of the vector of fitness effects of mutations *sj* across the genotype network. In other words, we use the correspondence between mutations at different genotypes to “transport” their fitness effects from one genotype to the other and finally to compare them. We emphasize that this interpretation is crucial for γ. Mathematically, the correlation of fitness effects can be defined only for some types of genotype spaces, like Hamming graphs, lattice graphs or more general Cartesian products of graphs. These are the only spaces where γ, which can always be defined by equation (14), could be interpreted as epistasis. This class includes all genotype spaces composed of multiple loci, multiple alleles at each locus and mutations acting on a single locus at a time, *i.e*. all spaces corresponding to DNA, RNA or protein sequences.

Correlations of fitness effects are not only useful to quantify epistasis. Their decay γ_d_ contains information on the nature of epistatic interactions and can reveal interesting signals. An example of that is the *Aspergillus* landscape studied here. The correlations γ_*d*_ for this landscape show an os-cillatory behaviour instead of the expected decay for random epistasis (*i.e*. HoC like) or for incompatibilities (*i.e*. Ising like), pointing towards a strong contribution of “eggbox-like” epistasis (reciprocal sign epistasis across multiple mutations). While the presence of pairwise reciprocal sign epistasis is not strange - it is actually quite common in compensatory interactions - the fact that reciprocal sign epistasis involves the whole landscape is quite surprising. In other words, starting from a first mutation chosen to be deleterious, it is not unreasonable that the second mutation could have a compensatory effect, but the mechanism behind the deleterious effect of the *third* mutation and the compensatory effect of the *fourth* mutation is obscure. It relate to complex pathways of interactions at the molecular level.

Many measures can only be computed if one has fitnesses for all combinations of the set of mutations (or subsets of). For example, the number of peaks lose meaning in a landscape with missing data, since the definition of a fitness maximum requires the knowledge of the fitness of all its neighbors. Since the fitness correlation functions *ρ*_*d*_ can be computed even with missing data, the correlation of fitness effects can be estimated from equation (14) even for very sparse landscapes. The sparseness of the landscape could increase the error on the estimate, however this effect could be compensated by the larger size of the landscape. Landscapes containing a larger number of mutations would be also more representative of real gene or protein landscapes.

For some landscapes, only fitness ranks or the beneficial/deleterious nature of the fitness effects can be experimentally determined. Our measure γ* is appropriate for these landscapes. While γ depends not only on positive and negative epistasis, but it is sensitive to its strength, γ* is based essentially on the fitness graph and therefore depends only on the sign of epistasis. γ and γ* are strongly correlated across fitness landscape models. Thus a mismatch between γ and γ* in real landscapes could point to some peculiar nature of epistatic interactions.

Finally, the γ and γ_*d*_ measures could also be useful to estimate parameters of theoretical landscape models from empirical data, thanks to the availability of approximate analytical formulae for these quantities. For example, assuming that the underlying model of a landscape is the NK model, the measures 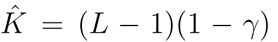 is an approximately unbiased Method-of-Moments estimator of the parameter *K*, *i.e*. 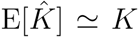 (see eq. 4). A similar approach can be used for the parameters of other landscape models. The potential of these measures for model inference and goodness-of-fit tests is yet to be studied.

We are still far from predicting evolution on real landscapes based on their measures, partly because of the incomplete knowledge of the structure of real landscapes, and partly because of the lack of measures with a natural evolutionary interpretation. In the future, we expect to witness a strong increase in the number of published empirical landscapes that will be experimentally resolved. The measures that we propose here will therefore find applications in the understanding and classification of these landscapes, as well as in studies of model landscapes. The correlation of fitness effects is a natural measure of epistasis that is comparable across landscapes, while the decay of correlations with mutation distance could be a useful tool to discriminate and classify these landscapes.

## Acknowledgments

We thank Joachim Krug, Ivan Szendro, Johannes Neidhart, Arjan de Visser, Nick Barton, RA Watson for useful comments and discussions. The work was funded by grant ANR-12-JSV7-0007 from Agence Nationale de la Recherche. BS acknowledges support from the Bonn-Cologne Graduate School.

## Appendix A. Common landscape measures

In this section we present some common landscape measures that can be applied as statistics for experimental landscape data. Notation: a genotype *g* is a sequence of alleles *g* = (*A*_1_*A*_2_*A*_3_… *A*_*L*_) of length *L*. For biallelic landscapes, *A*_*i*_ ∈ {0,1} and *S*_*i*_ = 2*A*_*i*_ − 1.

Some of the most common measures for fitness landscapes *ƒ*(*g*) are:

- number of peaks [32]: it is the number of genotypes such that all their neighbours have lower fitness, *i.e*. the number of local fitness maxima.
- *r/s* (roughness/slope) ratio [33]: the landscape is fitted to a linear model (a linear combination of *A*_*i*_s plus a constant) by least squares. The slope *s* is the average modulus of the coefficients of the *A*_*i*_s. The roughness *r* is the quadratic mean of the residuals of the regression. The measure of epistasis is their ratio *r/s*.
- fraction of epistatic interactions [34, 35]: the fraction of all pairs of mutations from all possible genotypes that show magnitude, sign or reciprocal sign epistasis.
- number of accessible paths [26]: assume that the absolute fitness maximum corresponds to the genotype g = (111…1). Count the number of paths of mutations 0 → 1 starting from g = (000…0) to (111…1) such that fitness increases after each mutation. This is the number of direct accessible paths to the maximum from its antipodal genotype.
- Fourier expansion and spectrum [36, 37, 31]: The coefficients a_*i*_1_…*i*_*J*__ of the Fourier expansion are uniquely defined in terms of the Fourier decomposition

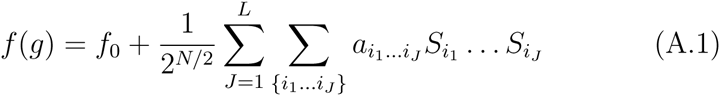

where {*i*_1_…*i*_*j*_} are ordered sets. The Fourier spectrum is defined by the sum of squared coefficients for interactions of *J* loci: 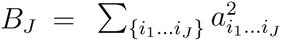. Epistasis is usually measured by the fraction of nonlinear interactions

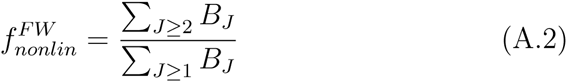

More details can be found in the review by Szendro et al. [23].

## Appendix B. Models of fitness landscapes

In this section we briefly illustrate some common models of fitness landscapes that will be used in this study. Most of them are illustrated in Figure B.5. Please note that we only considered here models of *L* biallelic loci. A mathematical formulation of these models is given in the main text.

**Figure B.5:**
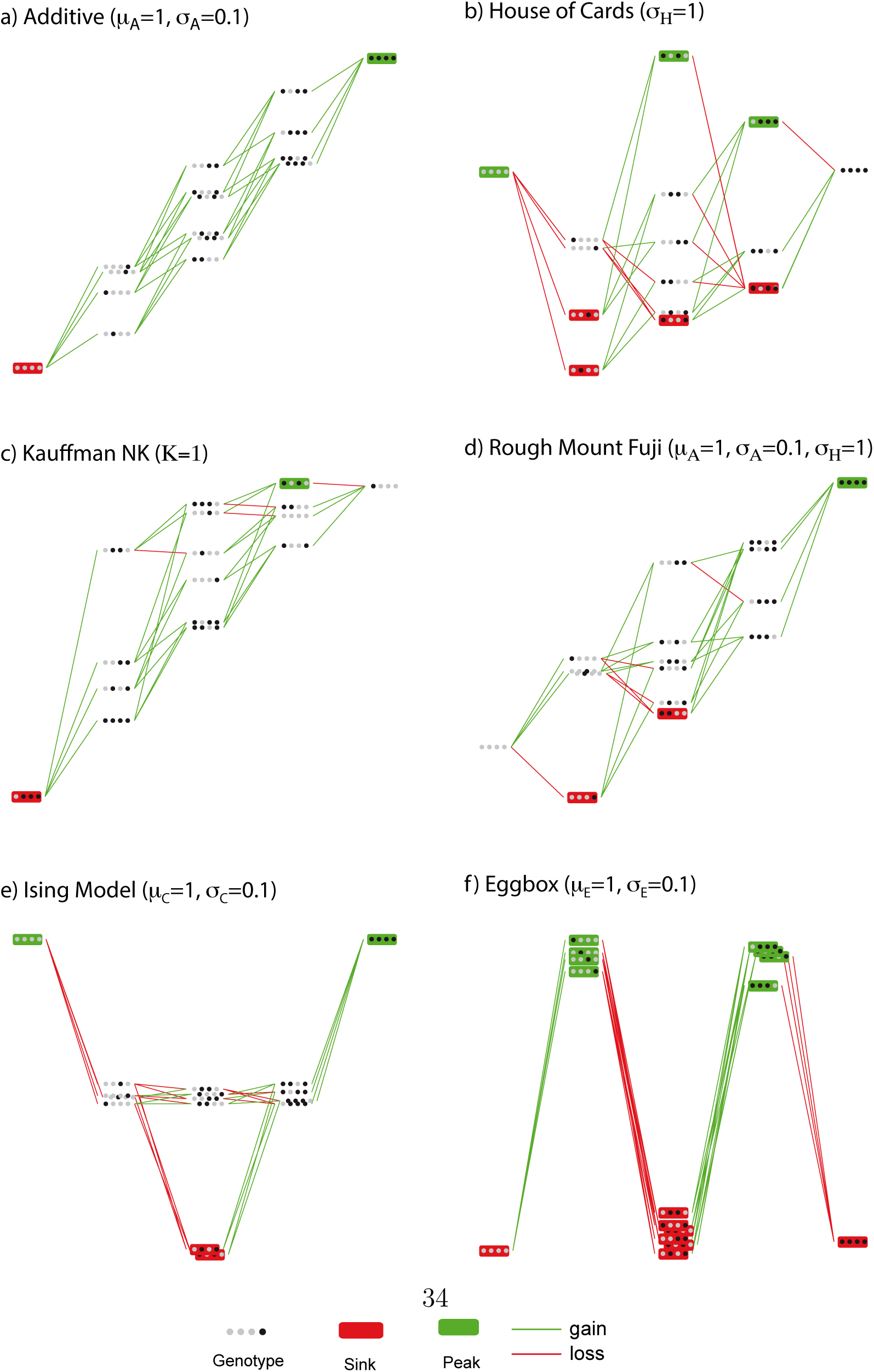
Models of fitness landscapes. Realizations of random landscapes obtained from the models discussed in the introduction, using Magellan [25].

*Appendix B.1. The Additive model (a.k.a. multiplicative model)*

This is a model for non-interacting mutations with independent fitness effects. The fitness is simply the product of the fitness contributions of each locus: fitness effects of different mutations are multiplied. In log-scale, this corresponds to summing the fitness effect of each mutation; for this reason this models is called “additive”. Here, the fitness effects of different mutations are randomly drawn from a Gaussian distribution with mean *μ*_*a*_ and variance 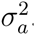. As there an independent contribution of each locus, the dimension of interaction is 1 (since each locus “interacts” only with itself).

In terms of Fourier decomposition, in this landscape all coefficients of second order and higher are zero.

*Appendix B.2. The House-of-Cards (HoC) model*

This is a model for random, uncorrelated fitness landscapes [18]. The fitness of each genotype is independent on the fitnesses of other genotypes. Here, it is randomly drawn according to a Gaussian of mean 0 and variance 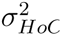. As this models corresponds to full interaction between the loci, the dimension of interaction is *L*.

In terms of Fourier decomposition, the coefficients are random variables with a marginal Gaussian distribution centered in 0.

*Appendix B.3. The Rough Mount Fuji (RMF) model*

This model interpolates between additive and uncorrelated fitness landscapes by adding the two [38]. The fitness is computed as the sum of an additive contribution and a HoC contribution. Here, the model is tuned by three parameters: mean *μ*_*a*_ and variance 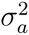 for the additive part and variance 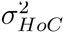 for the HoC part. (In the literature, this model is often defined with constant additive fitness effects, 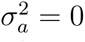). The model converges to an additive model when 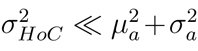 and to a HoC model when 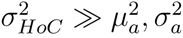. The dimension of interactions is a mixture of dimension 1 and dimension *L*.>

The Fourier decomposition is a linear function of the landscape, so it is a combination of the additive and the HoC decompositions.

*Appendix B.4. The NK model*

This landscape model with *N* = *L* loci interpolates between additive and uncorrelated fitness landscapes by combining uncorrelated fitness contributions (i.e. small HoCs) from *L* groups of *K* + 1 loci in an additive way [39]. There are different ways to choose the groups of interacting loci and while several properties such as mean number and mean height of local optima depend only weakly on the particular choice made [32], others seem to behave quantitatively different for some interaction choices [40]. Nonetheless it has been shown that the fitness correlation function is strictly independent of the interaction choice [24] and consequently γ does not depend on it either, while the number of chains may still be influenced by it. The number of interacting loci is *K* + 1 and the interpolation is controlled by the parameter *K* ∈ {0,1…*L* – 1}: *K* = 0 corresponds to an additive model with independent contributions from each locus, while *K* = *L* – 1 corresponds to an HoC model. The dimension of interaction is *K* + 1.

*Appendix B.5. The Ising, IMF and spin glass models*

This model originates from statistical physics [41], but has an immediate interpretation in terms of pairwise allele incompatibilities. In this model, each pair of interacting loci with different alleles causes a reduction in fitness. Here, loci interact only if they are neighbors in the genotype sequence (locus *i* interacts only with locus *i* – 1 and *i* + 1) and the first and the last locus have a single interaction (loci are arranged on a string). The fitness cost for each pair is drawn from a Gaussian with mean *μ*_*c*_ and variance 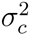. More general models based on allelic incompatibilities correspond to the Sherrington-Kirkpatrick model and other spin glass models in statistical physics [41]. The dimension of interaction is 2 as interactions only occurs between pairs. We also combined incompatibility interactions (Ising model) with an independent fitness contribution (additive model) in an “Ising Mount Fuji” (IMF) model in the same way the RMF is set.

In terms of Fourier decomposition, in this landscape all coefficients of third order and higher are zero.

*Appendix B.6. The Eggbox and the EMF models*

This model represents the extreme example of reciprocal sign epistasis of highest dimension. In this model, all genotypes in the landscapes have either low or high fitness. All the neighbours of a high-fitness genotype have low fitness, and vice versa. Therefore, in this landscape, each mutation is either deleterious (from high to low fitness) or compensatory (from low to high fitness). Fitnesses are given by a Gaussian with mean *ƒ*_0_ ± *μ*_*E*_/2. The dimension of interactions in this landscape is *L*. We also combine the eggbox interactions with independant contributions in an “Eggbox Mount Fuji” (EMF) landscape that is built like the RMF or the IMF.

In terms of Fourier decomposition, this landscape is dominated by the contribution of the Lth-order term (that is, the term of highest order).

## Appendix C. Proofs

*Appendix C.1. Relation between* γ *and type of epistasis*

The most extreme values of γ for different types of epistasis are obtained in the case *L* = 2. Denote by *ƒ*_00_, *ƒ*_10_, *ƒ*_01_, *ƒ*_11_ the log-fitness values. The function γ is defined as

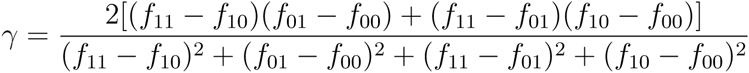

Since γ is a correlation, – 1 ≤ γ ≤ 1. γ is a continuous function and it is also invariant under permutations of loci and alleles, so from now on we will restrict to the subspace with *ƒ*_00_ ≤ *ƒ*_01_ ≤ *ƒ*_10_. Each of the three partitions of this subspace corresponding to magnitude, sign and reciprocal sign epistasis is connected, therefore the image of each one of them under γ is an interval.

By definition, magnitude epistasis results in γ ≥ 0 since all fitness jumps have the same sign. The extreme values 0 and 1 are both realized: γ = 0 in landscapes with *ƒ*_00_ = *ƒ*_01_ = *ƒ*_10_ < *ƒ*_11_, while γ = 1 in landscapes with *ƒ*_11_ = *ƒ*_10_ + *ƒ*_01_ – *ƒ*_00_. Therefore, for magnitude epistasis, 0 ≤ γ ≤ 1.

Reciprocal sign epistasis require that all fitness jumps change in sign after a mutation, therefore results in γ < 0 by definition. The extreme case γ = – 1 is realized in the landscape *ƒ*_01_ = *ƒ*_10_ > *ƒ*_00_ = *ƒ*_11_, while the other extreme case γ → 0 is realized for the landscapes with *ƒ*_00_ < *ƒ*_01_ = *ƒ*_10_ > *ƒ*_11_ for (*ƒ*_00_ – *ƒ*_01_) → 0. Therefore, for reciprocal sign epistasis, – 1 ≤ γ < 0.

Finally, sign epistasis can have both signs of γ. It is easy - although tedious - to show that there are no critical points of γ inside the space of landscapes with sign epistasis (with *L* = 2), therefore the extremal values should appear on the border. There are essentially two borders: (*ƒ*_11_ – *ƒ*_10_)→ 0 and *ƒ*_11_ = *ƒ*_01_. On the first border, we have γ > 0 and the upper limit is γ → 1 for the landscapes with *ƒ*_00_ = *ƒ*_01_ < *ƒ*_10_. On the second border, we have 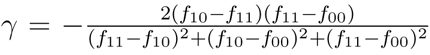 that reaches the minimum value γ = –1/3 at the landscapes with *ƒ*_10_ – *ƒ*_11_ = *ƒ*_11_ –*ƒ*_00_ (imposing the derivative with respect to *ƒ*_10_ to be null). Therefore, for sign epistasis, –1/3 ≤ γ < 1.

*Appendix C.2. Relation (14) between* γ_*d*_ *and the fitness correlation function*

Denote the fitness correlation function at distance *d* by *ρ*_*d*_ = Cor[*ƒ*(*g*), *ƒ*(*g*_*d*_)]. We use the identity

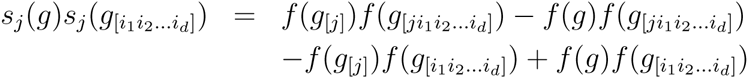

Averaging over the all mutations and all genotypes, the above terms give rise to:

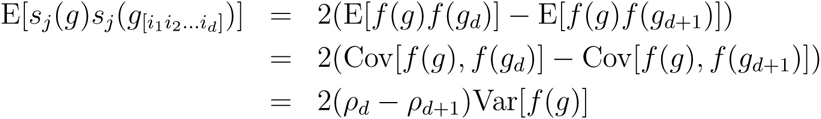

Summing over genotypes and mutations, the numerator of (13) becomes 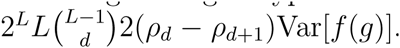. The denominator of (13) can be computed in a similar way by choosing *d* = 0, obtaining 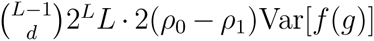. γ_*d*_ is their ratio

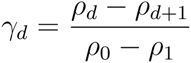

and since *ρ*_0_ = Cor[*ƒ*(*g*), *ƒ*(*g*)] = 1, we obtain the result (14).

*Appendix C.3. Relation (12) between* γ* *and the fractions of square motifs*

We consider a landscape without ties (*i.e*. all genotypes have different fitness values). In this case, 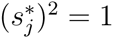 and therefore 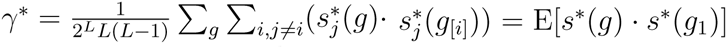 where the average is over all genotypes and pairs of mutations, or equivalently over all square motifs and over their sides. The average over the two sides of a motif is E[*s*\*(*g*) · *s*·(*g*_1_)] = (1 + 1)/2 = 1 with magnitude epistasis, (1 – 1)/2 = 0 with sign epistasis and (–1 – 1)/2 = –1 with reciprocal sign epistasis. To obtain the global average, we multiply these results by the fraction of motifs of each kind, *i.e*. γ* = 1 · *ϕ*_*m*_ + 0 · *ϕ*_*s*_ – 1 · *ϕ*_*rs*_. Since they sum to 1, we have *ϕ*_*m*_ = 1 – *ϕ*_*s*_ – *ϕ*_*rs*_ and substituting it we obtain the relation (12).

*Appendix C4. Relation (25) between* γ *and the Fourier spectrum*

We define 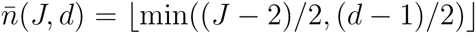. Since the Fourier basis is orthonormal, each component of the spectrum gives an independent contribution to the numerator and denominator of γ. The contribution of each *BJ* to the denominator of (3) is 4*J* since there are *J* nonzero mutations with fitness effect ±2a_*i*_1_…*i*_*J*__ each. For the numerator, there is again a factor *J* contributions (from nonzero mutations) multiplied by the square of the fitness effect of each term (averaged over the choice of the other *d* mutations). The fitness effect is ±4a_*i*_1_…*i*_*J*__ if an odd number of the *d* mutations lie within the indices *i*_1_…*i*_*J*_, and 0 otherwise. The probability that this number is odd is given by the sum of odd terms of the hypergeometric distribution with parameters *d*, *J* – 1,L – 1, therefore we have

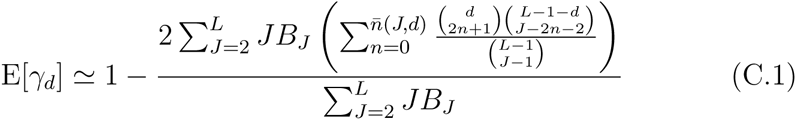

and since 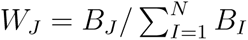, we have the equation (25).

## References

[1] S. Wright, The roles of mutation, inbreeding, crossbreeding and selection in evolution, Proc. 6th Int. Cong. Genet. 1 (1932) 356–366.

[2] H. A. Orr, The genetic theory of adaptation: a brief history, Nat Rev Genet 6 (2) (2005) 119–27. doi:10.1038/nrg1523.

[3] J. A. G. M. de Visser, J. Krug, Empirical fitness landscapes and the predictability of evolution, Nat Rev Genet 15 (7) (2014) 480–90. doi:10.1038/nrg3744.

[4] H. Richter, Fitness landscapes: From evolutionary biology to evolutionary computation, in: Recent Advances in the Theory and Application of Fitness Landscapes, Springer, 2014, pp. 3–31.

[5] D. L. Stein, Spin glasses and biology, Spin Glasses and Biology. Series: Series on Directions in Condensed Matter Physics, ISBN: 978-9971-5-0537-0. WORLD SCIENTIFIC, Edited by Daniel L Stein, vol. 6 6.

[6] J. H. Gillespie, A simple stochastic gene substitution model, Theor Popul Biol 23 (2) (1983) 202–15.

[7] S. Gavrilets, Fitness Landscapes and the Origin of Species, Princeton Uiversity Press, 2004.

[8] L.-M. Chevin, G. Decorzent, T. Lenormand, Niche dimensionality and the genetics of ecological speciation, Evolution 68 (5) (2014) 1244–56. doi:10.1111/evo.12346.

[9] F. A. Kondrashov, A. S. Kondrashov, Multidimensional epistasis and the disadvantage of sex, Proc Natl Acad Sci U S A 98 (21) (2001) 12089–92. doi:10.1073/pnas.211214298.

[10] J. A. G. M. de Visser, S.-C. Park, J. Krug, Exploring the effect of sex on empirical fitness landscapes, Am Nat 174 Suppl 1 (2009) S15–30. doi:10.1086/599081.

[11] S. P. Otto, The evolutionary enigma of sex, Am Nat 174 Suppl 1 (2009) S1–S14. doi:10.1086/599084.

[12] R. A. Watson, D. M. Weinreich, J. Wakeley, Genome structure and the benefit of sex, Evolution 65 (2) (2011) 523–36. doi:10.1111/j.1558-5646.2010.01144.x.

[13] S. Kauffman, The Origins of Order: Self-Organization and Selection in Evolution, Oxford University Press, 1993.

[14] N. Colegrave, A. Buckling, Microbial experiments on adaptive landscapes, Bioessays 27 (11) (2005) 1167–73. doi:10.1002/bies.20292.

[15] L.-M. Chevin, G. Martin, T. Lenormand, Fisher’s model and the genomics of adaptation: restricted pleiotropy, heterogenous mutation, and parallel evolution, Evolution 64 (11) (2010) 3213–31. doi:10.1111/j.1558-5646.2010.01058.x.

[16] M. L. M. Salverda, E. Dellus, F. A. Gorter, A. J. M. Debets, J. van der Oost, R. F. Hoekstra, D. S. Tawfik, J. A. G. M. de Visser, Initial mutations direct alternative pathways of protein evolution, PLoS Genet 7 (3) (2011) e1001321. doi:10.1371/journal.pgen.1001321.

[17] P. C. Phillips, Epistasis-the essential role of gene interactions in the structure and evolution of genetic systems, Nat Rev Genet 9 (11) (2008) 855–67. doi:10.1038/nrg2452.

[18] J. Kingman, A simple model for the balance between selection and mutation, Journal of Applied Probability (1978) 1–12.

[19] S. Kauffman, S. Levin, Towards a general theory of adaptive walks on rugged landscapes, J Theor Biol 128 (1) (1987) 11–45.

[20] J. Franke, A. Klözer, J. A. G. M. de Visser, J. Krug, Evolutionary accessibility of mutational pathways, PLoS Comput Biol 7 (8) (2011) e1002134. doi:10.1371/journal.pcbi.1002134.

[21] P. Hegarty, A. Martinsson, et al., On the existence of accessible paths in various models of fitness landscapes, The Annals of Applied Probability 24 (4) (2014) 1375–1395.

[22] J. Berestycki, E. Brunet, Z. Shi, The number of accessible paths in the hypercube, arXiv preprint arXiv:1304.0246.

[23] I. G. Szendro, M. F. Schenk, J. Franke, J. Krug, J. A. G. M. de Visser, Quantitative analyses of empirical fitness landscapes, Journal of Statistical Mechanics: Theory and Experiment.

[24] P. R. Campos, C. Adami, C. O. Wilke, Optimal adaptive performance and delocalization in NK fitness landscapes, Physica A: Statistical Mechanics and its Applications 304 (3) (2002) 495–506.

[25] S. Brouillet, H. Annony, L. Ferretti, G. Achaz, Magellan: a tool to visualize small fitness landscapes, biorXiv 031583, http://wwwabi.snv.jussieu.fr/public/Magellan/Magellan.main.html.

[26] D. M. Weinreich, N. F. Delaney, M. A. Depristo, D. L. Hartl, Darwinian evolution can follow only very few mutational paths to fitter proteins, Science 312 (5770) (2006) 111–4. doi:10.1126/science.1123539.

[27] J. A. de Visser, R. F. Hoekstra, H. van den Ende, Test of interaction between genetic markers that affect fitness in Aspergillus niger, Evolution 51 (1997) 1499–1505.

[28] J. A. Draghi, J. B. Plotkin, Selection biases the prevalence and type of epistasis along adaptive trajectories, Evolution 67 (11) (2013) 3120–31. doi:10.1111/evo.12192.

[29] D. Greene, K. Crona, The changing geometry of a fitness landscape along an adaptive walk, PLoS Comput Biol 10 (5) (2014) e1003520. doi:10.1371/journal.pcbi.1003520.

[30] F. Blanquart, G. Achaz, T. Bataillon, O. Tenaillon, Properties of selected mutations and genotypic landscapes under fisher’s geometric model, arXiv 1405.3504.

[31] J. Neidhart, I. G. Szendro, J. Krug, Exact results for amplitude spectra of fitness landscapes, J Theor Biol 332 (2013) 218–27. doi:10.1016/j.jtbi.2013.05.002.

[32] E. D. Weinberger, Local properties of Kauffman’s NK model: A tunably rugged energy landscape, Physical Review A 44 (10) (1991) 6399.

[33] T. Aita, M. Iwakura, Y. Husimi, A cross-section of the fitness landscape of dihydrofolate reductase, Protein Eng 14 (9) (2001) 633–8.

[34] D. M. Weinreich, R. A. Watson, L. Chao, Perspective: Sign epistasis and genetic constraint on evolutionary trajectories, Evolution 59 (6) (2005) 1165–74.

[35] F. J. Poelwijk, D. J. Kiviet, D. M. Weinreich, S. J. Tans, Empirical fitness landscapes reveal accessible evolutionary paths, Nature 445 (7126) (2007) 383–6. doi:10.1038/nature05451.

[36] P. F. Stadler, Landscapes and their correlation functions, Journal of Mathematical Chemistry 20 (1) (1996) 1–45.

[37] D. M. Weinreich, Y. Lan, C. S. Wylie, R. B. Heckendorn, Should evolutionary geneticists worry about higher-order epistasis?, Curr Opin Genet Dev 23 (6) (2013) 700–7. doi:10.1016/j.gde.2013.10.007.

[38] T. Aita, H. Uchiyama, T. Inaoka, M. Nakajima, T. Kokubo, Y. Husimi, Analysis of a local fitness landscape with a model of the rough mt. fuji-type landscape: application to prolyl endopeptidase and thermolysin, Biopolymers 54 (1) (2000) 64–79. doi:10.1002/(SICI)1097-0282(200007)54:1<64::AID-BIP70>3.0.CO;2-R.

[39] S. A. Kauffman, E. D. Weinberger, The NK model of rugged fitness landscapes and its application to maturation of the immune response, J Theor Biol 141 (2) (1989) 211–45.

[40] B. Schmiegelt, J. Krug, Evolutionary accessibility of modular fitness landscapes, Journal of Statistical Physics 154 (2014) 334–355.

[41] M. Mézard, M. A. Virasoro, G. Parisi, Spin glass theory and beyond, World scientific, 1987.

